# Identification of *SMARCD1* as a syndromic intellectual disability gene that is required for memory and context-dependent regulation of neuronal genes in *Drosophila*

**DOI:** 10.1101/422188

**Authors:** Kevin C.J. Nixon, Justine Rousseau, Max H. Stone, Mohammed Sarikahya, Sophie Ehresmann, Seiji Mizuno, Naomichi Matsumoto, Noriko Miyake, DDD study

**Affiliations:** Department of Physiology and Pharmacology, Schulich School of Medicine and Dentistry, Western University, London, Ontario, Canada; CHU-Sainte Justine Research Center, University of Montreal, Montreal, QC, Canada; Department of Biology, Faculty of Science, Western University, London, Ontario, Canada; Division of Genetics and Development, Children’s Health Research Institute, London, Ontario, Canada; Department of Pediatrics, Central Hospital, Aichi Human Service Center, Aichi, Japan; Yokohama City University Graduate School of Medicine, Yokohama 236-0004, Japan; Faculty of Medicine, University of Southampton, Southampton, UK; Belfast City Hospital, Belfast, Northern Ireland, UK; Perelman School of Medicine, University of Pennsylvania, Philadelphia, PA, USA; AP-HP, Hôpital Pitié-Salpêtrière, Département de Génétique et de Cytogénétique; Centre de Référence Déficience Intellectuelle de Causes Rares; GRC UPMC « Déficience Intellectuelle et Autisme », F-75013, Paris, France; INSERM, U 1127, CNRS UMR 7225, Sorbonne Universités, UPMC Univ Paris 06 UMR S 1127, Institut du Cerveau et de la Moelle épinière, ICM, F-75013, Paris, France; IGBMC, CNRS UMR 7104/INSERM U964/Université de Strasbourg, 67400 Illkirch, France; Institute of Human Genetics, University Hospital Essen, University of Duisburg-Essen, 45 239 Essen, Germany; Birmingham Women’s and Children’s NHS Foundation Trust Mindelsohn Way, Birmingham, UK; Department of Pediatrics, University of Montreal, Montreal, QC, Canada

**Author notes:** These authors contributed equally to this work (Kevin C.J. Nixon, Justine Rousseau). These authors contributed equally to this work; corresponding authors. Address: Philippe Campeau, Division of Medical Genetics, Room 6727, Sainte-Justine Hospital, 3175 Cote-Sainte-Catherine, Montreal QC Canada H3T 1C5, Jamie Kramer, Physiology and Pharmacology Medical Science Building, Room 266, Western University, 1151 Richmond St, London, Ontario, Canada, N6A 5C1.

## Abstract

Mutations in several genes encoding components of the SWI/SNF chromatin remodeling complex cause syndromic intellectual disability (ID). Here, we report on 5 individuals with mutations in the *SMARCD1* gene, presenting with ID, developmental delay, hypotonia, feeding difficulties, and small extremities. The mutations were proven to be *de novo* in 4 of the 5 individuals. Mutations in other SWI/SNF components cause Coffin-Siris, Nicolaides-Baraitser, or other syndromic ID disorders. Although the individuals presented here have some clinical overlap with these disorders, they lack the typical facial dysmorphisms. To gain insight into the function of SMARCD1 in neurons, we investigated the *Drosophila* ortholog, Bap60, in postmitotic memory-forming neurons of the adult *Drosophila* mushroom body (MB). Targeted knockdown of Bap60 in the MB of adult flies causes defects in long-term memory. Mushroom body specific transcriptome analysis revealed that Bap60 is required for context-dependent expression of genes involved in neuron function and development in juvenile flies when synaptic connections are actively being formed in response to experience. Taken together, we identify *SMARCD1* mutations as a novel cause of ID, and establish a role for the SMARCD1 ortholog Bap60 in regulation of neurodevelopmental genes during a critical time window of juvenile adult brain development that is essential in establishing neuronal circuits that are required for learning and memory.

## Introduction

Regulation of gene expression in neurons is critical for normal brain development and for normal cognitive functioning in adult animals^1–4^. Chromatin structure is an important factor in modulating gene expression^5^. Disruption of genes encoding chromatin regulators can result in intellectual disability (ID)^6,7^, a prevalent neurodevelopmental disorder characterized by limitations in adaptive behaviour and intellectual function. Mutations in several subunits of the SWI/SNF (SWItch/Sucrose Non-Fermenting) ATP-dependent chromatin remodeling complex (also known as the BAF complex in mammals) cause syndromic ID disorders, such as Nicolaides-Baraitser Syndrome (OMIM: 601358) and Coffin-Siris Syndrome (OMIM: 135900)^8–11^. SWI/SNF mutations are also potentially involved in other neurodevelopmental disorders such as schizophrenia, and autism spectrum disorder^12–15^, emphasizing the importance of this complex in neural development and function. In total, 9 of the 28 genes encoding SWI/SNF components have been implicated in ID, yet it remains to be determined if the remaining subunits are also involved in ID or other neuronal disorders.

The SWI/SNF complex is a highly conserved chromatin remodeling complex that hydrolyzes ATP to generate the energy required to alter nucleosome-DNA interactions resulting in more open chromatin for transcription factor binding and subsequent gene expression activation^3,16–18^. In mammals, there are multiple cell-type specific conformations of the SWI/SNF complex, including npBAF, specific for neuronal progenitors, and nBAF, specific for postmitotic neurons^1,19–22^. The SWI/SNF complex is important for regulation of gene expression programs involved in neuronal differentiation and brain-region specification in mice^2,4,19,23–27^. However, the complex is also essential in mature neurons for memory formation, synaptic plasticity, and activity responsive neurite outgrowth^3,4,28^.

Here, we characterize novel mutations in the *SMARCD1* gene in patients presenting with syndromic ID. *SMARCD1* encodes a core component of the SWI/SNF complex that has not previously been associated with ID, or any other neurodevelopmental condition. Furthermore, we show that the *Drosophila* SMARCD1 ortholog Bap60 is required in the mushroom body (MB) of adult flies for normal longterm memory. The MB is the learning and memory center of the fly brain^29,30^. We find that Bap60 has a profound effect on the expression of neurodevelopmental genes in the MB during a critical time window of juvenile brain development when synaptic connections are formed in response to early life experiences.

## Methods

### Patient enrolment

Individual 1 was enrolled in a study protocol that was approved by the institutional review boards of Yokohama City University School of Medicine. Individual 2 was enrolled in a study of the Groupe Hospitalier Pitié-Salpêtrière approved by the INSERM institutional review board. Individuals 3 and 4 were enrolled in the DDD study and in a study approved by the institutional review board of the CHU Sainte-Justine. Individual 5 had exome sequencing on a clinical basis and the family consented to the sharing of clinical information without photos. Genematcher was used to connect with the clinicians of individuals 2 and 5.

### Exome sequencing

For individual 1, genomic DNA was enriched for exons using the SureSelect All Human Exon kit (Agilent). Libraries were sequenced on the Illumina HiSeq, and analysis was performed as described^31^. For individual 2, exome sequencing was performed as described in^32^ (as for family 1 in that citation). For the individuals 3 and 4, exome sequencing was done as part of the DDD project. Exomes were enriched using the Agilent SureSelect 55MB Exome Plus library followed by Illumina HiSeq sequencing, and analysis was performed as described^33^. For individual 5, exome sequencing was done at GeneDx. Using genomic DNA from the proband and parents, the exonic regions and flanking splice junctions of the genome were captured using the Clinical Research Exome kit (Agilent). Sequencing was done on an Illumina system with 100bp or greater paired-end reads. Reads were aligned to human genome build GRCh37/UCSC hg19 and analyzed for sequence variants using a custom-developed analysis tool. Additional sequencing technology and variant interpretation protocol (e.g. Sanger) has been previously described^34^. The general assertion criteria for variant classification are publicly available on the GeneDx ClinVar submission page (http://www.ncbi.nlm.nih.gov/clinvar/submitters/26957/).

### Co-lmmunoprecipitation

HEK293T and SK-N-AS were cultured in DMEM, 10% FBS (Wisent), 1X AA and 1X Glutamax all from ThermoFisher Scientific. SK-N-AS media was supplemented with 0.1mM NEAA (ThermoFisher Scientific) and 25mM HEPES (Wisent). pcDNA.3.1-Flag-SMARCD1 wild type (NM_003076.4) or SMARCD1-c.990C>G (p.Asp330Glu), c.1457G>A (p.Trp486*) and c.1483T>C (p.Phe495Leu) were transfected using jetPRIME (Polyplus-transfection) according to the manufacturer’s instructions. After 48h, cells were washed once with ice-cold phosphate-buffered saline, lysed in a hypotonic buffer (25 mM HEPES pH 7.9, 25 mM KCL, 50mM EDTA, 5 mM MgCl2, 10% glycerol, 0.1% NP40, 1 mM DTT, complete mini protease inhibitors (Sigma-Aldrich)), and centrifuge at 3000 rpm for 3 minutes. The pelleted nuclei fraction (NE) was resuspended in nuclear lysis buffer (25 mM Tris, pH8, 150 mM NaCl, 1 mM EDTA, 1% triton, 5% glycerol, complete mini protease inhibitors), incubated 30 min at 4 degrees and centrifuge at 13 000 rpm for 20 min. For immunoprecipitation, nuclear extracts (100 to 300ug) were incubated with either pre-coupled M2 anti-FLAG (M8823, Sigma-Aldrich) or anti-Brg1 (H10) coupled to Dynabeads (ThermoFisher Scientific) magnetic beads. After 1.5 and 4 hours respectively, beads were washed 4 times and proteins eluted directly in 1X laemmli loading buffer supplemented with 50 mM DTT. Western blot was performed using the following antibodies: anti-SMARCC1 (PCRP-SMARCC1-1F1, DSHB), BRG-1 (H10) (sc-374197, Santa-Cruz), anti-FLAG M2 (368791, Sigma-Alrich), anti-BRG/BRM (J1) (a generous gift from J.Lessard, IRIC).

### Fly stocks and culture

Flies were reared on standard cornmeal-agar media with a 12h/12h light cycle in 70% humidity. The following stocks were obtained from the Bloomington *Drosophila* Stock Center (Indiana University): short hairpin Bap60 RNAi lines generated by the Transgenic RNAi Project (TRiP)^35^ (*UAS-Bap60^32503^* and *UAS-Bap60^33954^*); the TRiP short hairpin *mCherry^RNAi^* line (stock #35785); the mushroom body specific Gal4 driver line *R14H06-Gal4* (stock #48667) from the Janelia Flylight^36^ collection, and the temperature sensitive Gal80 (*Gal80^ts^*) driven by the tubulin promoter (stock #7019). *UAS-unc84::GFP* was a gift from G.L. Henry.

### Courtship Conditioning

Courtship conditioning assays were performed as described previously^37^. Training was performed by pairing individual males with premated wild type females. Long-term memory was induced through a seven-hour training period and tested 24 hours after training. For each fly pair, a courtship index (CI) was determined by manually observing the percentage of time spent courting during a 10-minute period. Statistically, loss of memory was identified using two complimentary methods. Reduction of the mean CI of trained (CI_trained_) flies compared to naïve (CI_naive_) of the same genotype was compared using a Kruskal-Wallis test followed by pairwise comparisons using Dunn’s test. A learning index (LI) was also calculated (LI=(CI_naive_-CI_trained_)/CI_naive_). LIs were compared between genotypes using a randomization test^52^(10,000 bootstrap replicates) performed with a custom R script^37^ and the resulting p-values were corrected for multiple testing using the method of Bonferroni.

The temperature-sensitive Gal80^ts^ system^38^ was used to specifically knockdown Bap60 in the adult mushroom body. Flies of the genotype *tubGal80^ts^; R14H06-Gal4* were crossed to Bap60 RNAi lines, *UAS-Bap60^32503^* and *UAS-Bap60^33954^*, as well as a control *UAS-mCherry^RNAi^* that is present in the same genetic background as the RNAi transgenes. Crosses were raised at 18ºC, the permissive temperature which allows Gal80^ts^ to repress Gal4 mediated transcription. At eclosion, male flies of the genotypes *Gal80^ts/+^; R14H06-Gal4/UAS-Bap60^32503^*, *Gal80^ts/+^; R14H06-Gal4/UAS-Bap60^33954^ and Gal80^ts/+^; R14H06-Gal4/UAS-mCherry^RNAi^* were transferred to 29ºC, which causes Gal80 inactivation, allowing Gal4 mediated induction of UAS-RNAi transgenes. After five days, collected males were tested for long-term memory using courtship conditioning as described above.

### Isolation of nuclei in tagged cell types (INTACT)

Fifty juvenile (0-3h old) or mature (1-5 days old) adult male flies of the genotype UAS-unc84::GFP/+; R14H06-Gal4/UAS-Bap60^32503^ (Bap60-KD) and UAS-unc84::GFP/+;R14H06-Gal4/UAS-mCherry^RNAi^ (control) were anesthetized using CO_2_ and flash frozen using liquid nitrogen. Protein G Dynabeads (Invitrogen) were adsorbed to 5μg of anti-GFP antibody (Invitrogen, G10362) in PBS 0.1% Tween-20 (PBST) for 10 minutes at room temperature. Beads were then isolated using a magnet and resuspended in PBST. The flies were then vortexed and placed in ice-cold sieves to separate and isolate their heads. Heads were then added to homogenization buffer (25mM KCl pH 7.8, 5mM MgCl2, 20mM Tricine, 150nM spermine, 500nM spermidine, 10nM β-glycerophosphate, 250mM sucrose, 1X Pierce protease inhibitor tablets – EDTA-free (ThermoFisher Scientific) or 1X Halt protease inhibitor cocktail – EDTA-free (ThermoFisher Scientific)), and the suspension was homogenized for approximately 1 minute using a standard tissue homogenizer at 30,000 rpm. The suspension was then placed in a Dounce homogenizer with NP40 (ThermoFisher Scientific) added to an end concentration of 0.3% and homogenized 6 times with the tight pestle. The homogenate was filtered using a 40μm strainer and pre-cleared with non-antibody bound beads for 10 minutes. Antibody-bound beads were added to the homogenate for 30 minutes, followed by washing in homogenization buffer. Total RNA was isolated from immunoprecipitated nuclei using the Arcturus PicoPure RNA isolation kit (ThermoFisher Scientific) with DNase digestion using the RNase-free DNase kit (Quiagen) according to the manufacturer’s instructions. Isolated RNA quality was then assessed using the Bioanalyzer 2100 Pico RNA kit (Agilent), by visual examination of rRNA peak integrity.

### RNA-sequencing and analysis

RNA from MB-enriched samples was used for RNA-seq library generation using the NuGEN Ovation *Drosophila* RNA-Seq System (BioLynx), according to the manufacturer’s instructions. Size selection using Agencourt SPRI select beads (Beckman-Coulter) was used to select for library sizes of 200bp. Library size was assessed using the Bioanalyzer 2100 DNA high sensitivity kit (Agilent). Sequencing was performed using the Illumina NextSeq500 at the London Regional Genomics Centre (Robarts Research Institute, London, Ontario) using the high output v2 75 cycle kit to a read length of 75bp for single-end reads.

Raw sequencing reads were trimmed using Prinseq quality trimming^39^ and a minimum base quality score of 30. Read quality was then assessed using FastQC (http://www.bioinformatics.babraham.ac.uk/projects/fastqc). Trimmed reads were aligned to the *Drosophila melanogaster* reference genome (BDGP release 6)^40,41^ using the STAR aligner^42^. An average of 54,803,904 and 46,899,292 high quality uniquely aligned reads with a maximum of four mismatches were obtained from juvenile Bap60-KD MBs and controls respectively and an average of 26,865,090 and 35,504,102 high quality uniquely aligned reads with a maximum of four mismatches were obtained from mature Bap60-KD MBs and controls respectively (**Table S1**). The number of reads per gene was quantified by inputting resulting reads into HTSeq-count^43^ with the --type flag indicating ‘gene’ (**Table S1**). Genes encoding rRNA and genes with no counts across all samples were removed from analysis, leaving 12,922 and 13,440 genes for downstream analysis for juvenile and mature samples, respectively.

Raw gene counts were normalized and differential expression analysis between Bap60-KD MBs and controls was performed using the R^44^ package DESeq2^45^. Differentially expressed genes were defined as genes with a fold-change >1.5 or > 2 and a Benjamini-Hochberg adjusted p-value <0.05. Gene ontology (GO) enrichment analysis was performed on upregulated and downregulated differentially expressed genes using Panther^46^ (FDR<0.05).

### Classification of tissue specific genes

To generate lists of tissue-specific genes, we used normalized gene expression values from Brown *et al.^47^* for several tissues including, adult head (9 samples), adult carcass (3 samples), and adult digestive tract (3 samples). The relative enrichment in gene expression levels for each of these tissues was calculated by comparing the mean bases per kilobase per million reads (bpkm) for each specific tissue to the mean bpkm across all remaining tissues. Tissue-specific and tissue-depleted genes are defined as having an average log2 fold-change in expression outside of one standard deviation of the fold-change in expression of all genes in all tissue types (**Table S2**). Adult heads were used to represent a neuron-enriched tissue, while the adult carcass, consisting of tissues remaining after removal of the head, digestive tract, and reproductive organs, represents a muscle-enriched tissue. A list of MB-specific genes (**Table S2**) was generated using published MB-specific gene expression that were obtained using the same INTACT protocol described here^48^. MB-specific genes were defined as genes that were significantly upregulated more than 2-fold in MB-enriched samples compared to biologically paired whole head input samples. Statistical significance of over or under representation of tissue-specific genes in differentially upregulated and downregulated genes in Bap60-KD MB flies was determined using a hypergeometric test.

## Results

### Human Genetics and Clinical Data

In a cohort of individuals collected because of clinical overlap with Coffin-Siris syndrome, *de novo* mutations in several genes encoding members of the SWI-SNF complex were identified^8^. As an expansion of this study, an individual was identified with a *SMARCD1* variant (p.Asp330Glu), but the *de novo* status of the variant could not be confirmed since the father was not available for testing. As part of the DDD study, two individuals were identified with *SMARCD1* variants (p.Trp486* and p.Phe495Leu). Through Genematcher, two additional individuals were identified, a patient with a *de novo* p.Arg446Gly variant in a cohort of individuals with agenesis of the corpus callosum, and another individual who had clinical exome sequencing for developmental delay and other anomalies who was identified to have a truncating variant (pArg503*).

The variants and the associated clinical features are shown in **Figure 1, Table 1, and Table S3**. There is phenotypic overlap with Coffin-Siris syndrome in that two patients have a hypoplastic 5^th^ toenail and all have ID, however, most do not have the typical facial features (wide mouth with thick everted lips, broad nose, thick eyebrows, and long eyelashes) (see **Figure 2** for pictures). All had feeding difficulties, and three had hypotonia. All individuals had developmental delay and none had epilepsy. Three individuals had small hands and feet. The dysmorphisms were variable and were most notable in individual 4 with the p.Phe495Leu variant, who also had the most profound neurodevelopmental disorder (at 3 years of age, he could not talk or walk).

**Figure 1:**
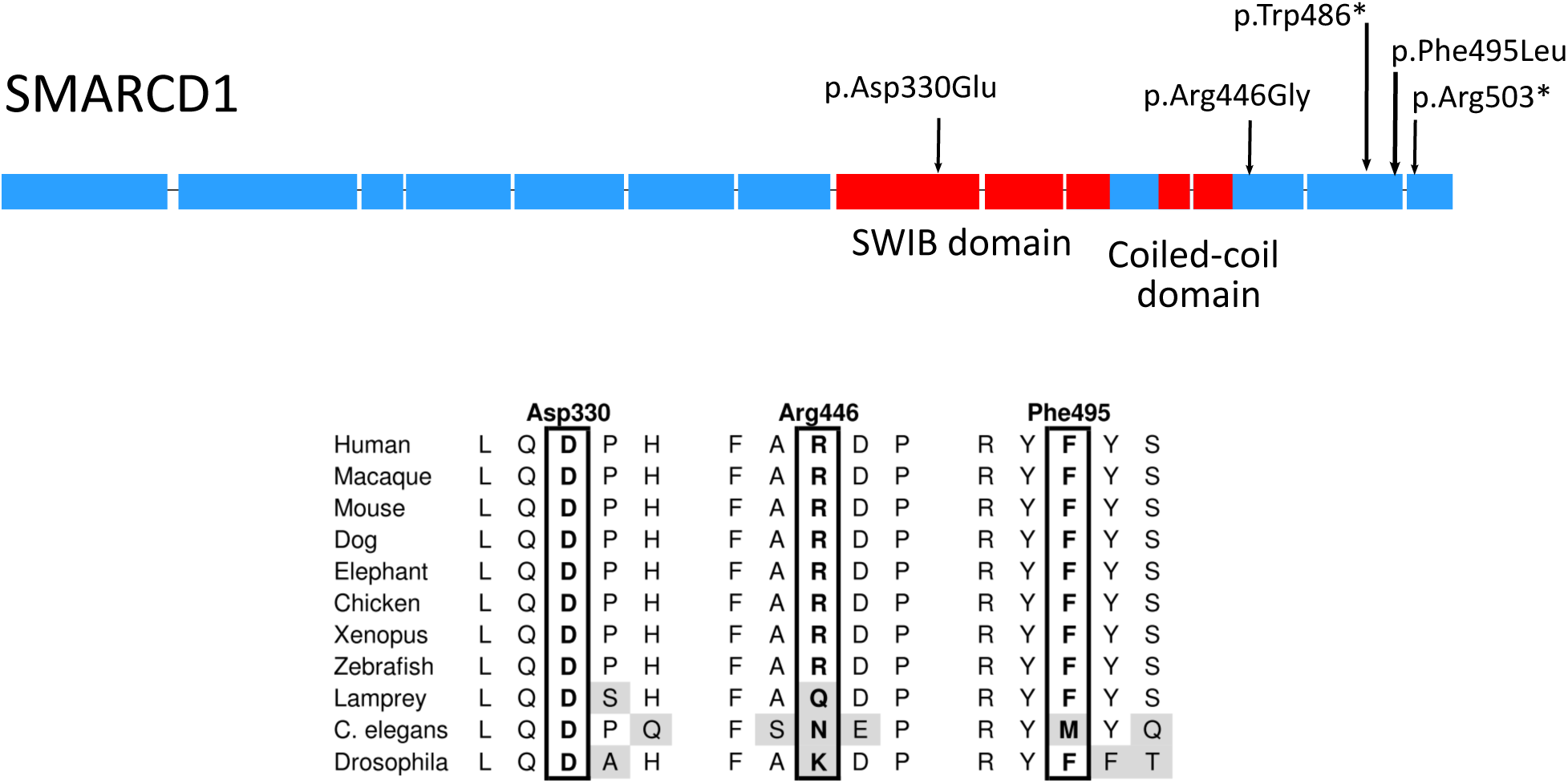
Protein structure of *SMARCD1* and associated domains with respective variants. SWIB and coiled-coil domains are shown in red. The location of the identified variants areindicated with black arrows. Conserved regions of *SMARCD1* across species are shown below.

**Table 1:**
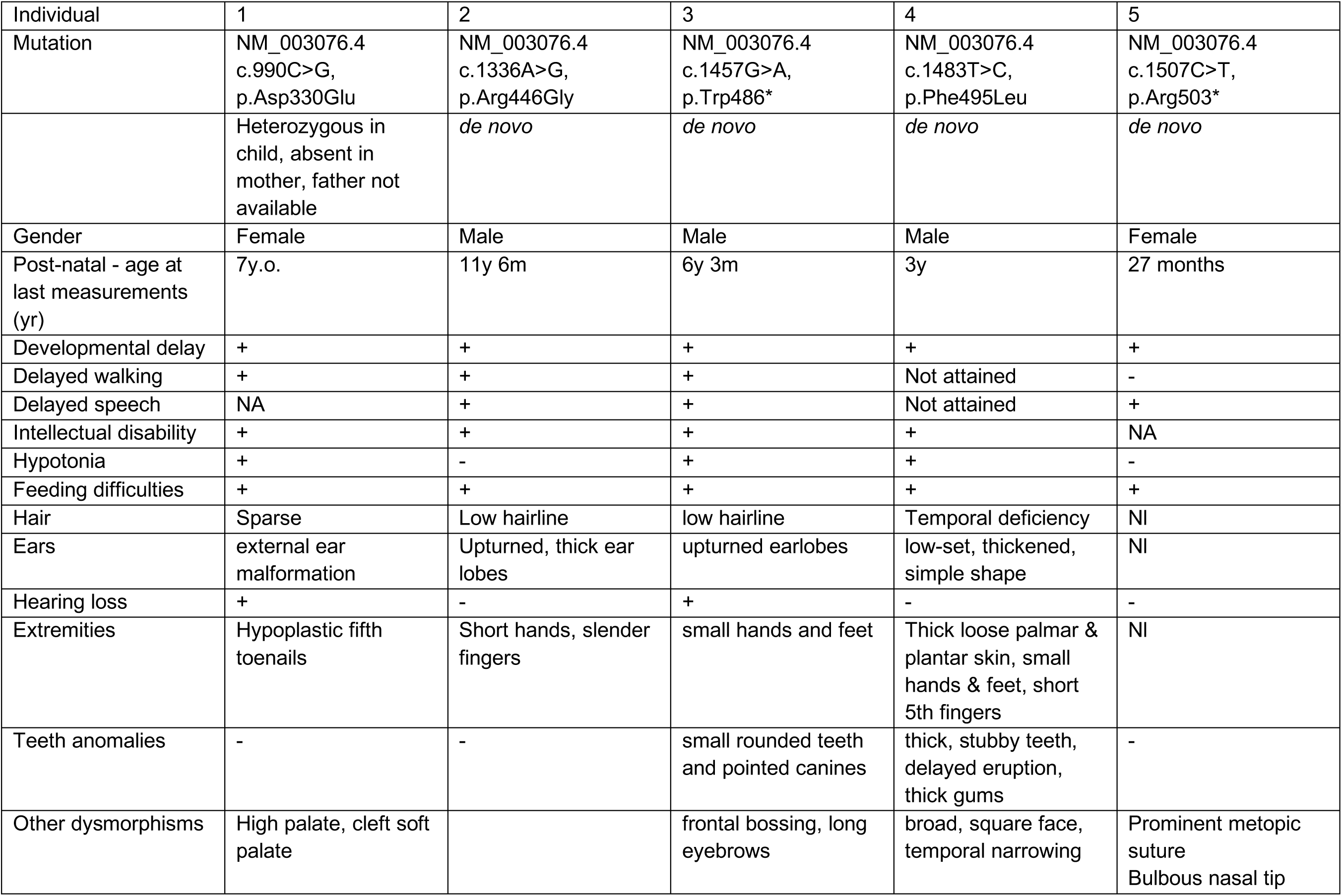
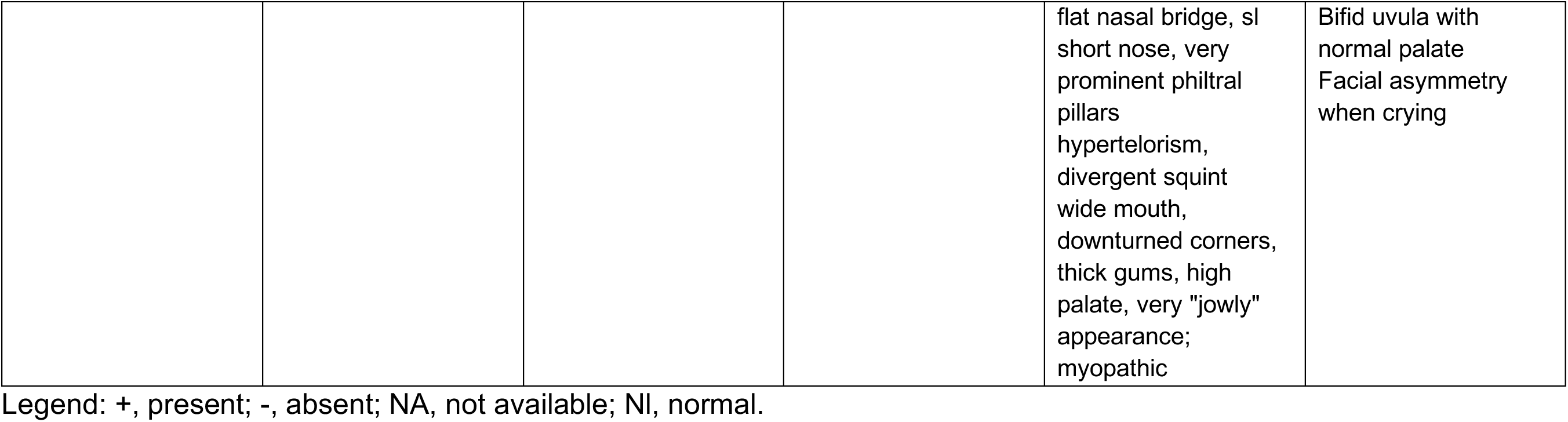
Summary of clinical findings (see Table S3 for additional details)

**Figure 2:**
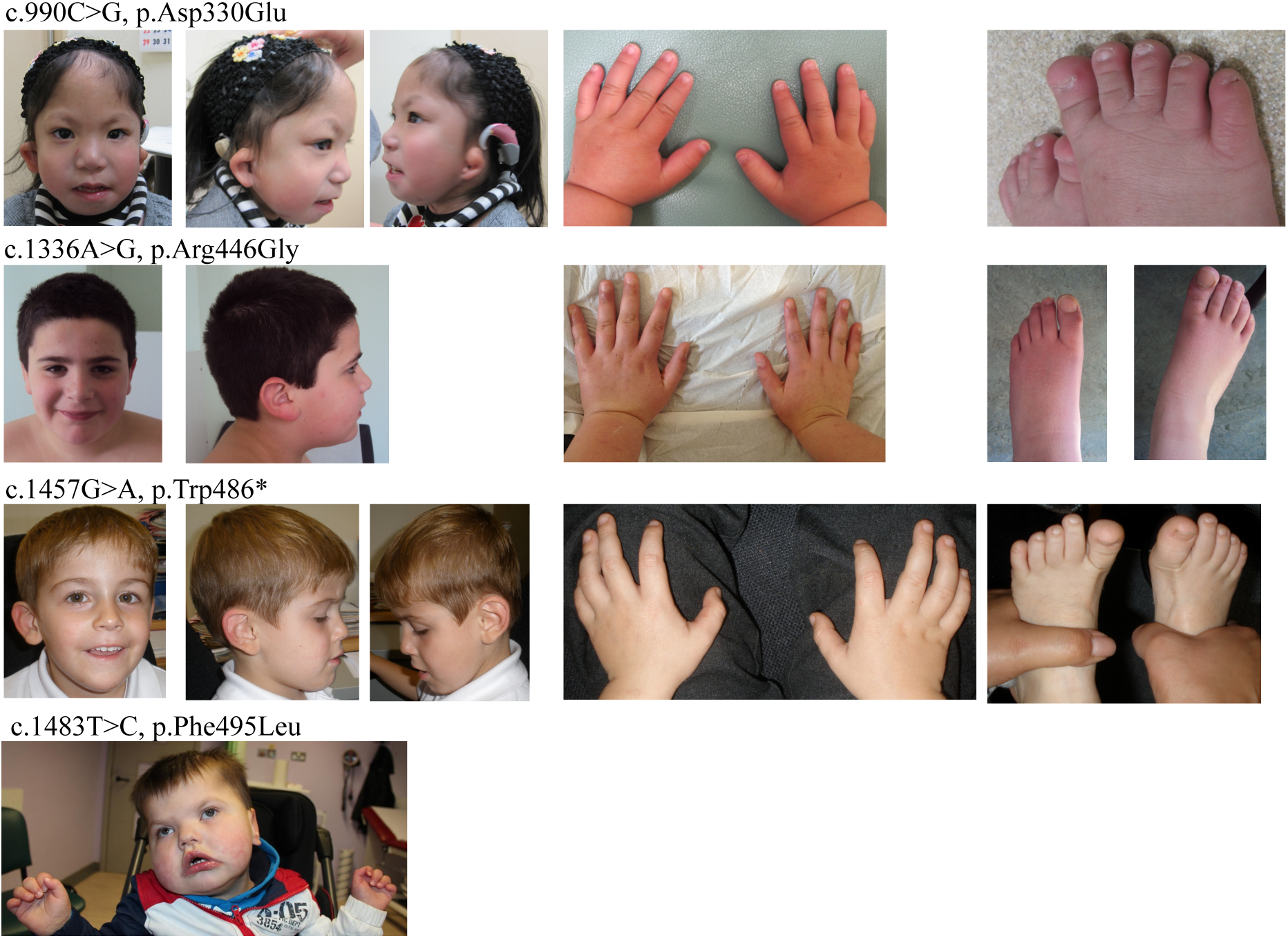
Photos of individuals identified with *SMARCD1* mutations.

### *In silico* assessment of the variants

We performed some bioinformatic analysis of the observed variants. *SMARCD1* has an ExAC pLI score of 1 (since only one loss-of-function (LoF) variant was observed in the ExAC cohort while 26 were predicted), indicating that the gene is intolerant to heterozygous LoF variants^49^. It also had a positive Exac Z-score of 3.95, indicating it is also relatively intolerant to missense variants (74 observed while 183 were predicted). DECIPHER gives a %HI score of 16.64%, which is close to the ranks indicating there is a high likelihood that the gene will exhibit haploinsufficiency (below 10%)^50^. DOMINO gives at 99.8% probability for the gene to be associated with an autosomal dominant condition^51^.

We used wANNOVAR to predict pathogenicity scores, those mostly considering conservation (polyphen, LRT, MutationAssessor, GERP, PhyloP, phastCons, SiPhy), function (SIFT, PROVEAN), or combination thereof (FATHMM, MutationTaster, DANN, CADD). We considered the following prediction cutoffs: damaging, rank scores above 0.5, and CADD phred scores above 25 (**Table S4**). Using these cutoffs, MutationTaster, FATHMM-MKL, and PROVEAN considered all assessed variants deleterious. LRT, GERP, PhyloP, phastCons, SyPhy, SIFT, DANN and CADD predicted most assessed variants to be deleterious. MutationAssessor gave medium scores, and polyphen2 predicted 2 out of 3 assessed variants to be benign. p.Arg446Gly was the variant with the most tools giving low pathogenicity predictions, mostly because of the poor conservation in animals further removed from humans (see *lamprey, C. elegans* and *Drosophila*, **Figure 1**).

### *In vitro* assessment of the variants

We generated expression constructs with three different variants (p.Asp330Glu, p.Trp486*, p.Phe495Leu). None of the mutations affect protein stability, as all were detectable at normal levels by Western blot. Upon co-immunoprecipitation, we noted that all three mutants bind well to BRG1 and SMARCC1 (**Figure S1**). Therefore, the mutations might not affect integration in the SWI/SNF complex, but they might affect the function of SMARCD1 otherwise, possibly by changing the activity of the complex or interactions with other proteins.

### The Drosophila SMARCD1 ortholog Bap60 is required for memory

Given that some of the mutations, such as the truncating mutations and the SWIB domain mutation might lead to a loss-of-function, we assessed in *Drosophila* the consequence of loss of Bap60, the Drosophila ortholog of SMARCD1. To do this, we generated Bap60 knockdown flies using previously characterized transgenic RNAi lines^52^. RNAi knockdown of Bap60 was targeted to the MB, the learning and memory center of the fly brain, using *R14H06-Gal4*. Knockdown was confined to adult flies using the temperature sensitive Gal4 inhibitor, Gal80^ts^ (see methods). We tested memory in MB-specific Bap60 knockdown flies using a classic paradigm known as courtship conditioning^37,53^. In this assay, male flies display a learned reduction of courtship behavior after sexual rejection by a non-receptive pre-mated female. Control flies expressing an RNAi construct targeting mCherry showed a normal reduction in courtship behavior after being sexually rejected by a mated female (**Figure 3A**). reduction in courtship was not seen in flies expressing RNAi transgenes targeting Bap60 (**Figure 3A**) and Bap60 knockdown flies had a significant reduction in the learning index (LI) compared to the controls (**Figure 3B**). Both Bap60 RNAi lines, which have unique target sequences, showed a similar reduction in memory, which suggests that memory defects are not due to off-target effects. These results suggest that Bap60 is required for normal memory, post development, in adult neurons.

**Figure 3:**
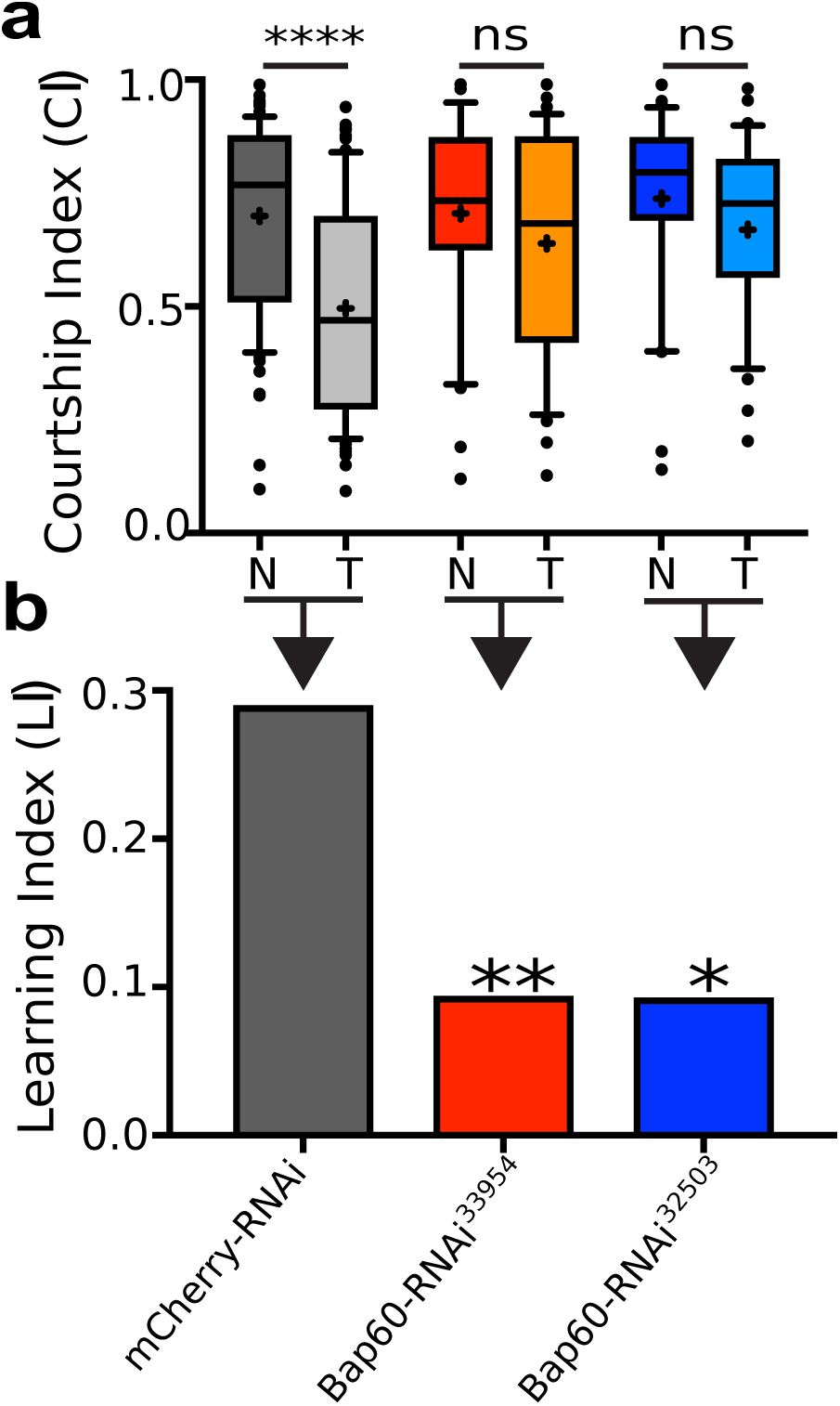
Adult-specific knockdown of Bap60 in the MB results in reduced capacity for both short-term and long-term courtship memory. (**A**) Boxplots showing courtship indices (CI) for the mCherry RNAi control and Bap60 knock down flies in long-term courtship conditioning assays (+ indicates the mean). The mean CI for naïve (N) and trained (T) flies of the same genotype was compared using a Wilcoxon test. (**B**) Learning indices (LI) for control and Bap60 knockdown flies. Adult specific Bap60 knockdown resulted in a significantly reduced LI (randomization test, 10,000 bootstrap replicates) relative to control flies.

### Bap60 is required for expression of neuron-specific genes during a critical period of juvenile adult MB development

We used INTACT (see methods) to investigate the effect of Bap60 knockdown on gene expression in the specific MB cells that were targeted for RNAi knockdown in our learning and memory assays. In juvenile adult insects, the mushroom body undergoes a period of development and synaptogenesis in the first hours after eclosion^54–60^. During this time, neuronal connections are formed that are critical for normal learning and memory later in life^56,58^. We compared the MB-specific transcriptome in Bap60 knockdown flies compared to controls in early juvenile adults (0-3 h after eclosion), and more mature adults (1-5 days after eclosion). We observed a greater effect of Bap60 on gene expression at the juvenile stage than in the mature adult MB (**Figure 4A and 4B**). Differential expression analysis between the juvenile Bap60-KD MBs and control using DESeq2^45^ yielded 416 differentially expressed genes (P_adj_<0.05 and Fold-Change > 1.5), of which 156 were upregulated and 260 were downregulated in the Bap60-KD MBs (**Figure 4A**, **Table S5**). In contrast, only 68 differentially expressed genes (29 upregulated and 39 downregulated) were observed in mature Bap60-KD MBs (**Figure 3B, Table S5**). Differential expression of several genes was confirmed by RT-qPCR in independent biological replicates (**Figure S2**). We performed gene ontology (GO) enrichment analysis to investigate the functions of differentially expressed genes. DE genes from the mature MB showed very little GO enrichment (**Table S5**). GO enrichment analysis of the upregulated genes from the juvenile MB revealed many terms related to muscle, such as “myofibril assembly” and “sarcomere organization” (**Table S5**). GO enrichment analysis of downregulated genes revealed neuron-related terms such as “neurotransmitter metabolic process”, “synaptic vesicle”, and “regulation of synaptic plasticity”, as well as developmental terms such as “nervous system development” and “anatomical structure development” (**Table S5**). This suggested an important role for Bap60 in the regulation of neurodevelopmental genes in the early juvenile adult MB.

Having observed a downregulation of neuron-related genes and an upregulation of muscle-related genes in Bap60-KD MBs, we reasoned that Bap60 may be required at this stage to activate expression of neuron-specific genes that contribute to cell identity. To test this, we used existing tissue specific RNA-seq data^47,48^ to establish a list of 904 “neuron-specific genes” that are enriched in heads compared to other tissues, and 118 “MB-specific genes” that are enriched in MB-specific INTACT samples compared to whole head (see methods, **Table S2)**. Of the 260 genes that are downregulated in Bap60-KD MBs 78 are neuron-specific, and 31 are MB-specific, which is significantly more than would be expected by random (p<10^-25^, hypergeometric test) (**Figure 4C**). On average these genes are expressed at a consistent level in controls in juvenile and mature adult MBs but have reduced expression in Bap60-KD MBs at the juvenile stage that recovers to normal levels in mature adults (**Figure 4D**). These trends were validated by RT-qPCR for a selection of genes in an independent experiment (**Figure S2**-*prt* and *jdp*). Muscle-specific genes (enriched in carcass compared to other tissues) were not significantly represented among genes that were downregulated in Bap60-KD MBs. Taken together, this data suggests Bap60 plays a context dependent role in activating the expression of neuron-specific genes in the MB.

**Figure 4:**
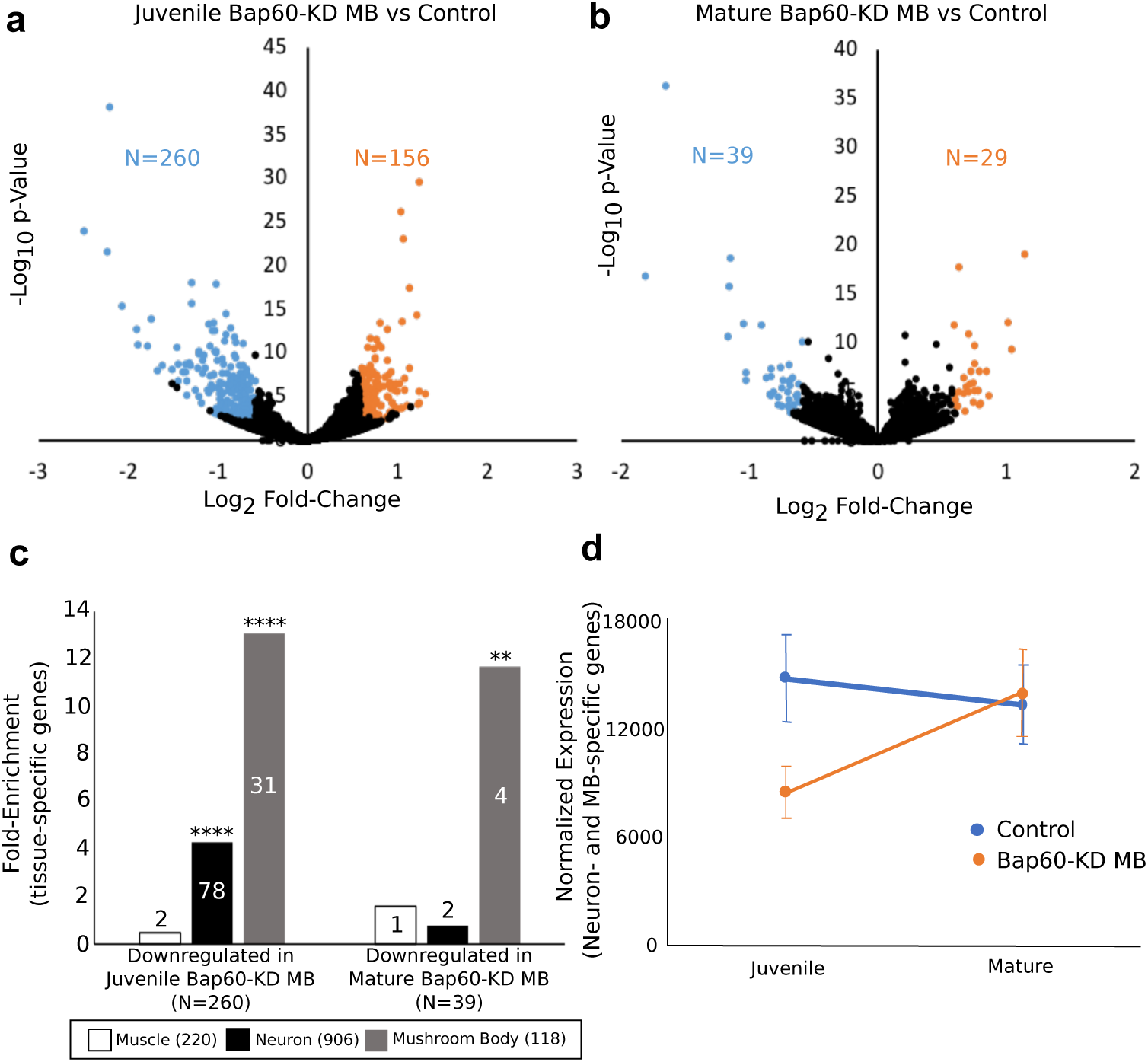
Mushroom body knockdown of Bap60 expression leads to dysregulation of tissue-specific genes. (**A** and **B**) Volcano plots of gene expression and p-Value. Differentially expressed genes (P_adj_<0.05 and fold-change > 1.5) are represented in blue (downregulated) or orange (upregulated) in (**A**) juvenile Bap60-KD MB and (**B**) mature Bap60-KD MB flies compared to controls of the same age. (**C**) Fold enrichment of muscle-, neuron-, and mushroom body-specific genes (Brown *et al.* and Jones *et al.* ^47,48^; see methods) in downregulated genes in juvenile and mature Bap60-KD MBs compared to the entire gene set analyzed (^**^P>10-3; ^****^P>10-25; Hypergeometric test). (**D**) Average normalized expression (±SEM)of neuron- and mushroom body-specific genes in control (blue) and *Bap60-KD MB* (orange) flies at the juvenile and mature stages.

### Bap60 is required for expression of developmental genes that are preferentially activated during juvenile MB development

In addition to neuronal genes, our GO enrichment analysis of genes that are downregulated in Bap60 mutants revealed many terms related to development (**Table S5**). We reasoned that Bap60 might be required for activation of key genes that are involved in experience dependent MB development in juvenile adults^58,61^. To test this, we performed DE analysis comparing the MB-specific transcriptome in early juvenile adults to mature adults (**Table S6**). In controls, 549 genes were significantly increased by 2-fold or more in the juvenile adult MB compared to mature adults. Of these 549 juvenile enriched MB genes, 385 genes were also > 2-fold enriched in Bap60-KD flies, while 165 were not (**Figure 5A**). On average, these 165 genes show significantly lower expression in juvenile Bap60-KD MBs compared to controls, and this difference is no longer observed in mature flies (**Figure 5B**). These expression trends were validated for a selection of genes in an independent RT-qPCR experiment (**Figure S2**). Interestingly, these 165 genes show a strong GO enrichment for terms related to development, and immune response (**Figure 5C**). In contrast, the 385 genes that show juvenile enrichment in both Bap60-KD and control MBs show very little GO enrichment. This suggests that Bap60 is required for activation of developmental genes during a critical period when the juvenile adult MB needs to establish new experience-dependent synaptic connections that are required for normal learning and memory throughout life^54–60^.

**Figure 5:**
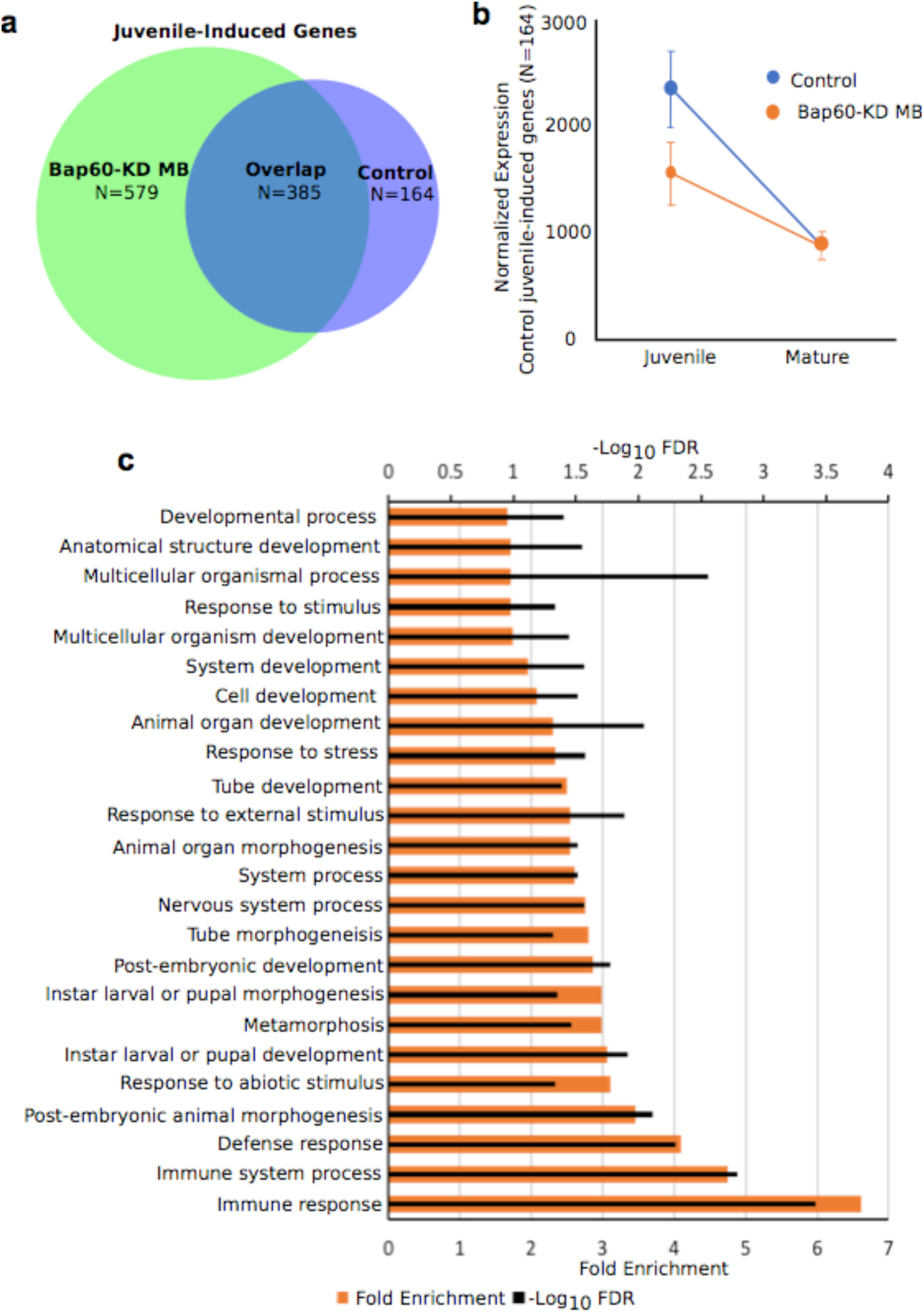
Mushroom body knockdown of Bap60 expression leads to dysregulation of developmental genes at the Juvenile stage. (**A**)Venn diagram showing overlap of upregulated genes (P_adj_<0.05 and fold-change > 2) between control and Bap60-KD MB files at the juvenile stage compared to a mature stage (juvenile-induced genes). (**B**)Average normalized expression (±SEM) of the 164 control juvenile-induced genes from (**A**) in both control (blue) and Bap60-KD MB (orange) flies at the juvenile and mature stages.(**C**) Gene ontology enrichment for biological processes of juvenile-induced genes control flies. Terms with at least 10 genes and an FDR < 0.05 are displayed.

## Discussion

### Characterization of a novel neurodevelopmental disorder

In this study, we describe a novel genetic disorder characterized by mutations in *SMARCD1*, which encodes a component of the SWI/SNF chromatin remodeling complex. Several other SWI/SNF genes are implicated in syndromic neurodevelopmental disorders that are typically characterized by ID, abnormalities of the 5^th^ digit, and characteristic facial features. The 5 individuals described here do fit in this spectrum, with ID and a low penetrance of 5^th^ digit abnormalities, but lack the typical facial dysmorphisms seen in other SWI/SNF related disorders (**Figure 2**, **Table 1**).

The identified *SMARCD1* variants, 2 nonsense and 3 missense, are all clustered in the C-terminal end of the protein (**Figure 1**). The two nonsense variants are located in the last exon of the transcript or within 50 base pairs of the last exon junction, so should escape nonsense mediated decay and produce a truncated protein. One missense variant is located in the highly conserved SWIB domain, while the other two are not located in any known domain. Although these missense and protein-truncating mutations are predicted to be damaging, we do not know the precise functional effect of these mutations. We show here that the proteins are stable and do not disrupt interaction with other members of the SWI/SNF complex, BRG1 and SMARCC1. These mutations might affect binding to other proteins such as transcription factors or other SWI/SNF components that we did not test. It is possible that these mutated proteins have a loss of function or dominant negative effect by integrating into the complex and compromising its normal activity, however, this remains to be tested. Notably, missense mutations are quite common in other SWI/SNF related disorders. While nonsense mutations are common in some SWI/SNF genes (*ARID1A, ARID1B,* and *ARID2*), *SMARCA4, SMARCA2, SMARCB1,* and *SMARCE1* are primarily affected by missense mutations with a predicted dominant negative effect^62^.

### Bap60 in memory and MB-specific transcriptome regulation in juvenile *Drosophila*

Although the role of some SWI/SNF components in neuron development and function is well described^63^, there was previously no work investigating the function of SMARCD1 in the nervous system. We show here that the *Drosophila* ortholog Bap60 is required in the adult fly mushroom body for normal memory (**Figure 3**). The requirement for Bap60 in the adult fly brain is consistent with evidence from mammals suggesting the presence of a SWI/SNF complex that is only present in differentiated neurons^3,28,64^. Indeed, it has been shown that the neuron-specific subunit BAF53b is also essential in adults for normal memory^3^. Taken together, this suggests that SWI/SNF-related ID might result from defective gene regulation postnatally in differentiated neurons.

Using MB-specific transcriptome analysis we found that Bap60 has a greater effect on gene regulation in the MB of juvenile adult flies compared to mature flies. In particular, Bap60 seems to be important for activating expression of neuronal genes (**Figure 4**) and developmental genes that normally show increased expression in juvenile MBs (**Figure 5**). This is interesting because the MB is known to undergo structural alterations and form new synaptic connections during the early stages of juvenile adult life^55,56,65,66^. These changes are dependent on sensory input, suggesting that some of the brain’s circuitry is developed in response to early life experience^54,55,66^. This early experience dependent plasticity in the mushroom body is required for normal memory at later life stages in flies^55,56,59,60^ and bees ^57,66^. Although much more complex, human brains also show periods of experience dependent plasticity, especially during adolescence^67^. So called “environmental enrichment” therapy for autism is based on the idea that defects in neural circuitry might be corrected by providing increased sensorimotor experience^67,68^. It will be interesting to see if SMARCD1 is required for experience-dependent brain plasticity in mammals.

## Acknowledgements

Thanks to the individuals and their families who participated in this study. We thank the Bloomington Drosophila Stock Center and the Transgenic RNAi Project at Harvard Medical School for creating and providing fly stocks used in this study. Thanks to David Carter at the London Regional Genomics Centre for help with sequencing. This work was funded by Canadian Institute of Health Research, the Canada Research Chairs program, the Canadian Foundation for Innovation, the Fonds de Recherche Santé Québec, and by AMED under grant numbers, JP18ek0109280, JP18dm0107090, JP18ek0109301, JP18ek0109348 and JP18kk020500; JSPS KAKENHI Grant Numbers, JP17H01539 and JP16H05160; and Takeda Science Foundation (N. Miyake, N. Matsumoto). We thank Agnès Rastetter for technical assistance and confirmation of the mutation proband 2. The DDD study presents independent research commissioned by the Health Innovation Challenge Fund (grant number HICF-1009-003), a parallel funding partnership between the Wellcome Trust and the Department of Health, and the Wellcome Trust Sanger Institute (grant number WT098051). The views expressed in this publication are those of the author(s) and not necessarily those of the Wellcome Trust or the Department of Health.

## Figures

**Figure S1:**
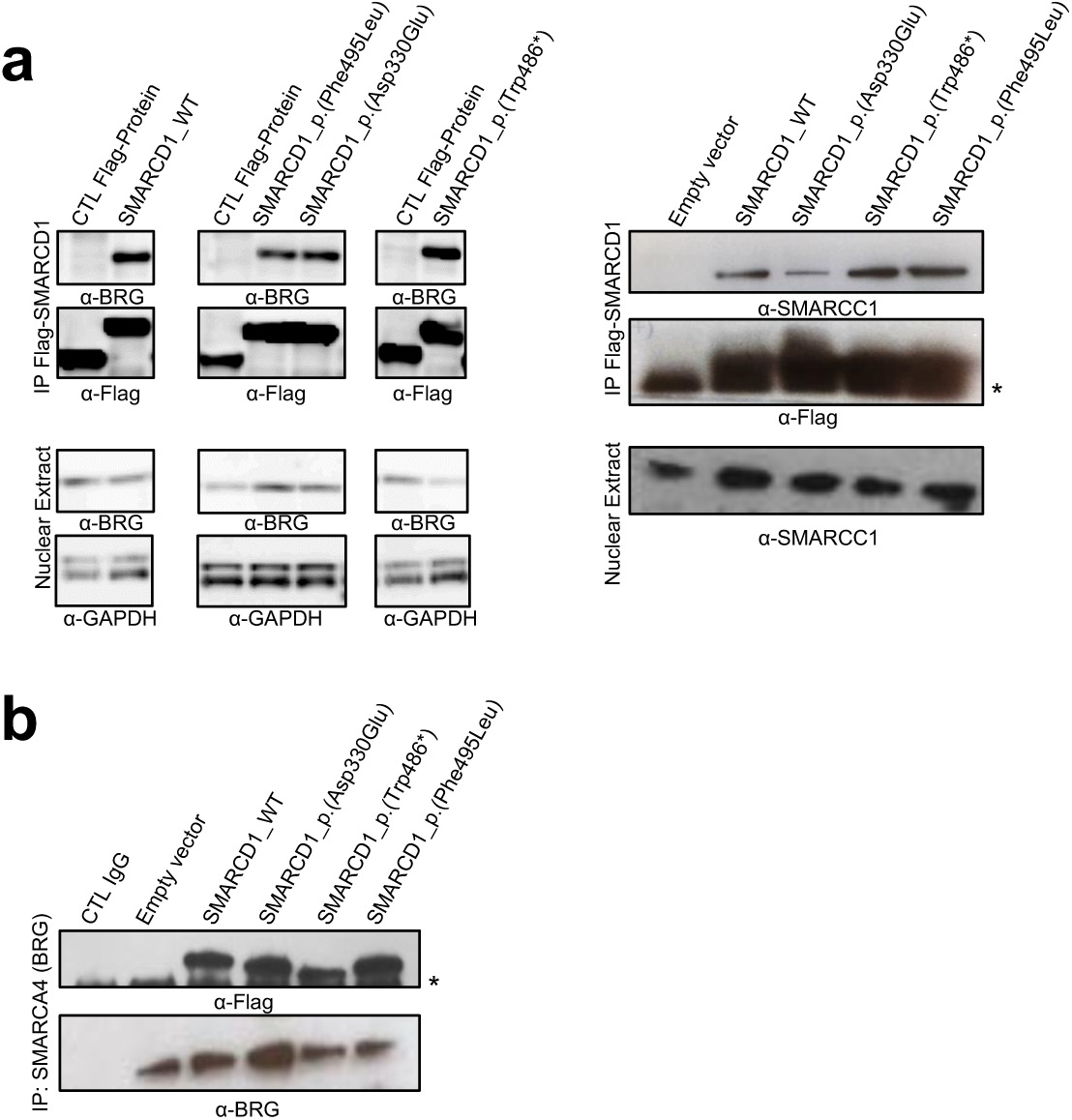
Mutations in SMARCD1 do not impair its binding to other SWI/SNF components. Co-Immunoprecipitation of SMARCA4 (BRG1) and SMARCC1 **(A)** and reciprocally of SMARCD1 **(B)** following immunoprecipitation of FLAG-SMARCD1 or endogenous SMARCA4 (BRG1) respectively.

**Figure S2:**
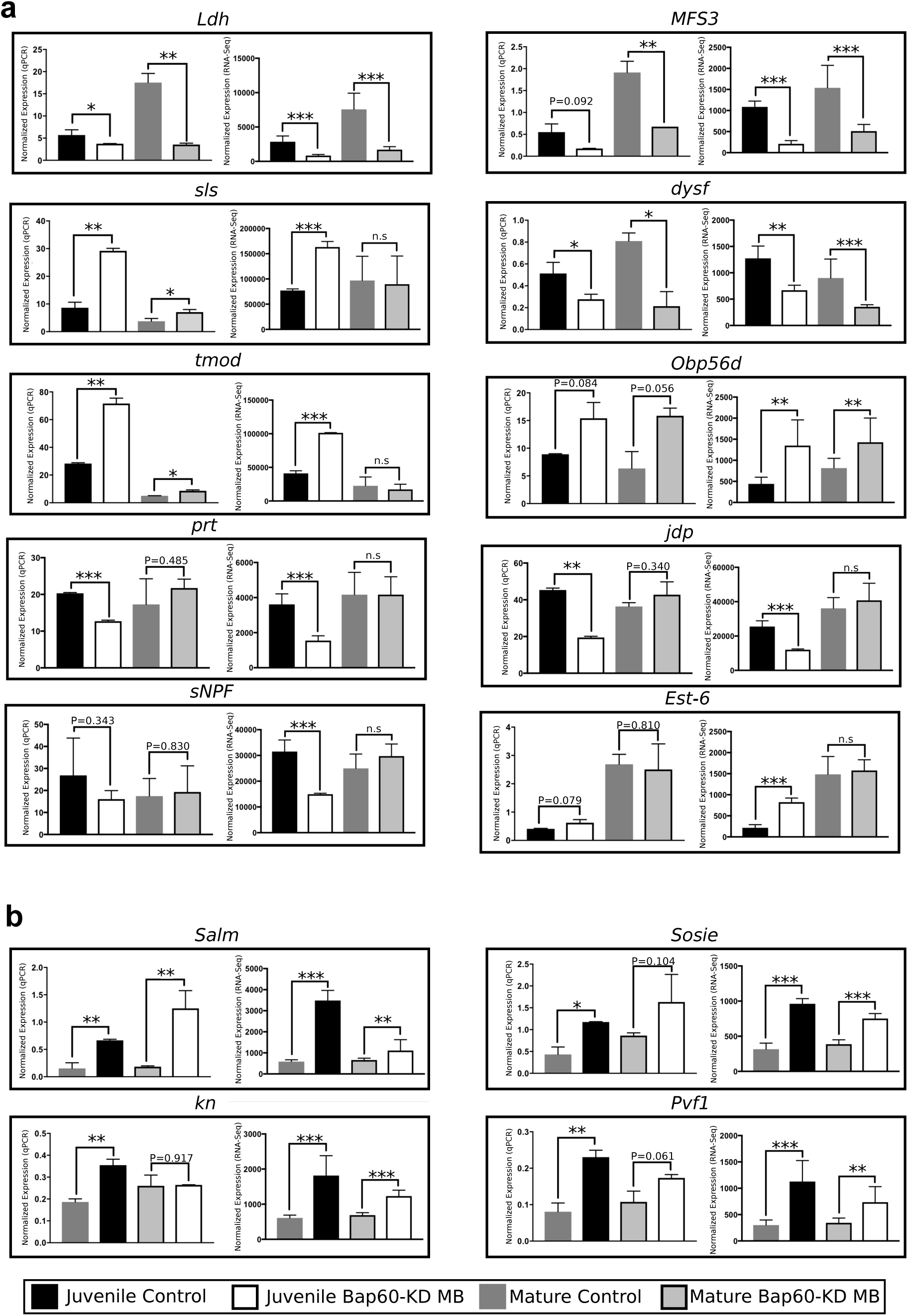
qPCR validation of RNAseq gene expression changes. Average normalized gene expression ± SD of genes as measured through RT-qPCR (left panel) and RNA-Seq (right panel) in Bap60-KD MB and control mushroom body (MB) samples. RT-qPCR gene expression is normalized to the housekeeping genes *Eif2β* and *βcop*and RNA-Seq gene expression is normalized from DESeq2. (**A**) Validation of overall gene expression trends for *Ldh, MFS3, sls, dysf, tmod, Obp56d, prt, jdp, sNPF*, and *Est-6.*> Comparisons were made between juvenile control and juvenile Bap60-KD MB and between mature control and mature Bap60-KD MB. Genes validated by RT-qPCR were: *Ldh, sls, dysf, tmod, prt*, and *jdp.* Genes that were showing similar trends between RT-qPCR and RNA-Seq were: *MFS3, Obp56d, and Est-6. sNPF* could not be validated by RT-qPCR due to variation between biological replicates. (**B**) Validations of gene expression trends of developmental genes: *Salm, Sosie, kn*, and *Pvf1* that were induced in juvenile control flies, but not juvenile Bap60-KD MB. Comparisons were made between mature control and juvenile control and between mature Bap60-KD MB and juvenile Bap60-KD MB. Genes validated were: *Kn* and *Pvf1. Sosie* is validated despite variability of the data. Not validated is: *Salm.* Significance for RT-qPCR is determined by Student’s t-test; significance for RNA-Seq is determined by binomial Wald test (DESeq2); ^*^P<0.05, ^**^P<0.01, ^***^P<0.001.

**Table S1.** Alignment summary of sequencing reads to the reference genome and counts to genic features using HTSeq-Count.

**Table S2.** Lists of neuron-specific and mushroom body-specific genes

**Table S3:**
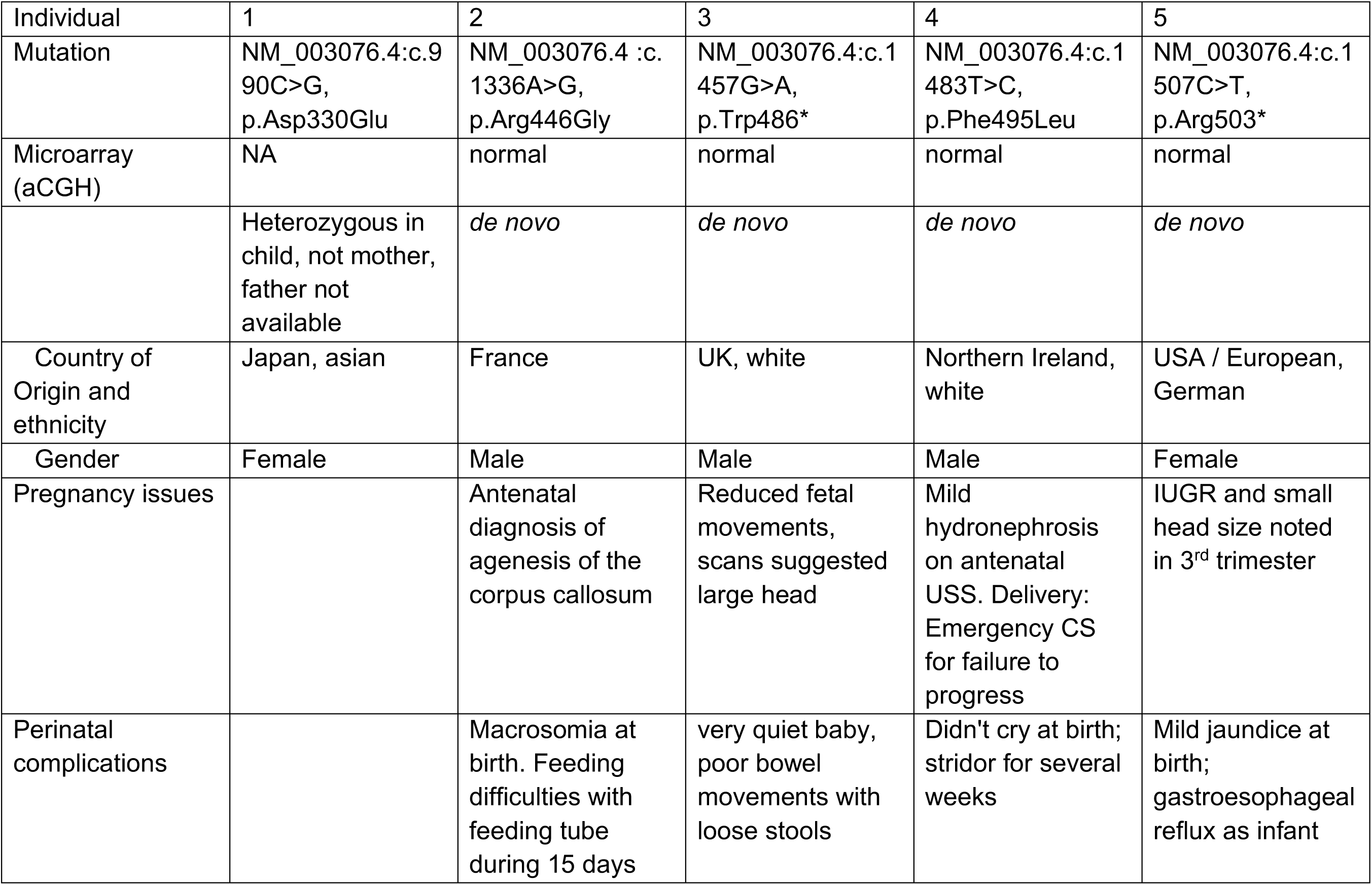
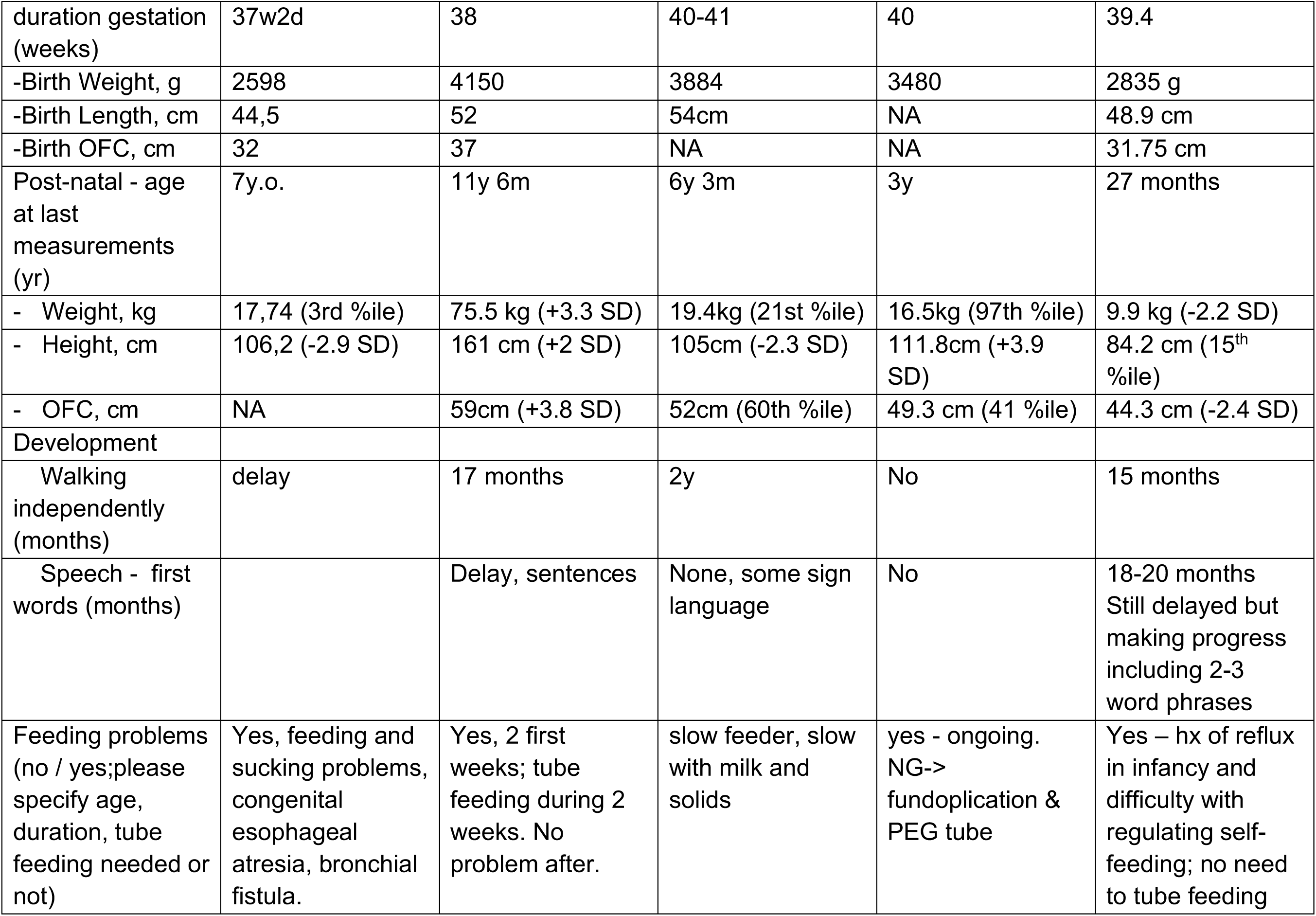
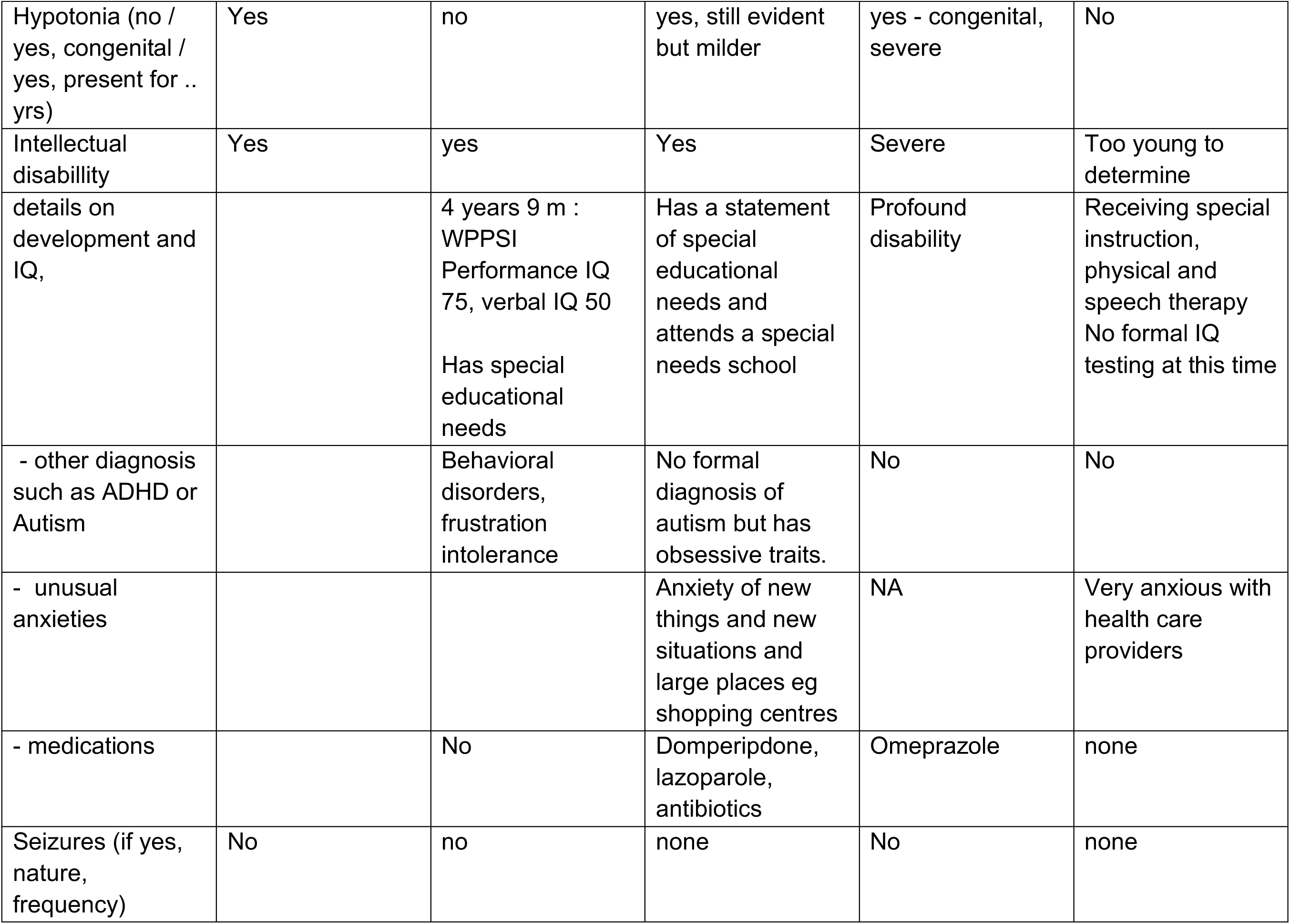
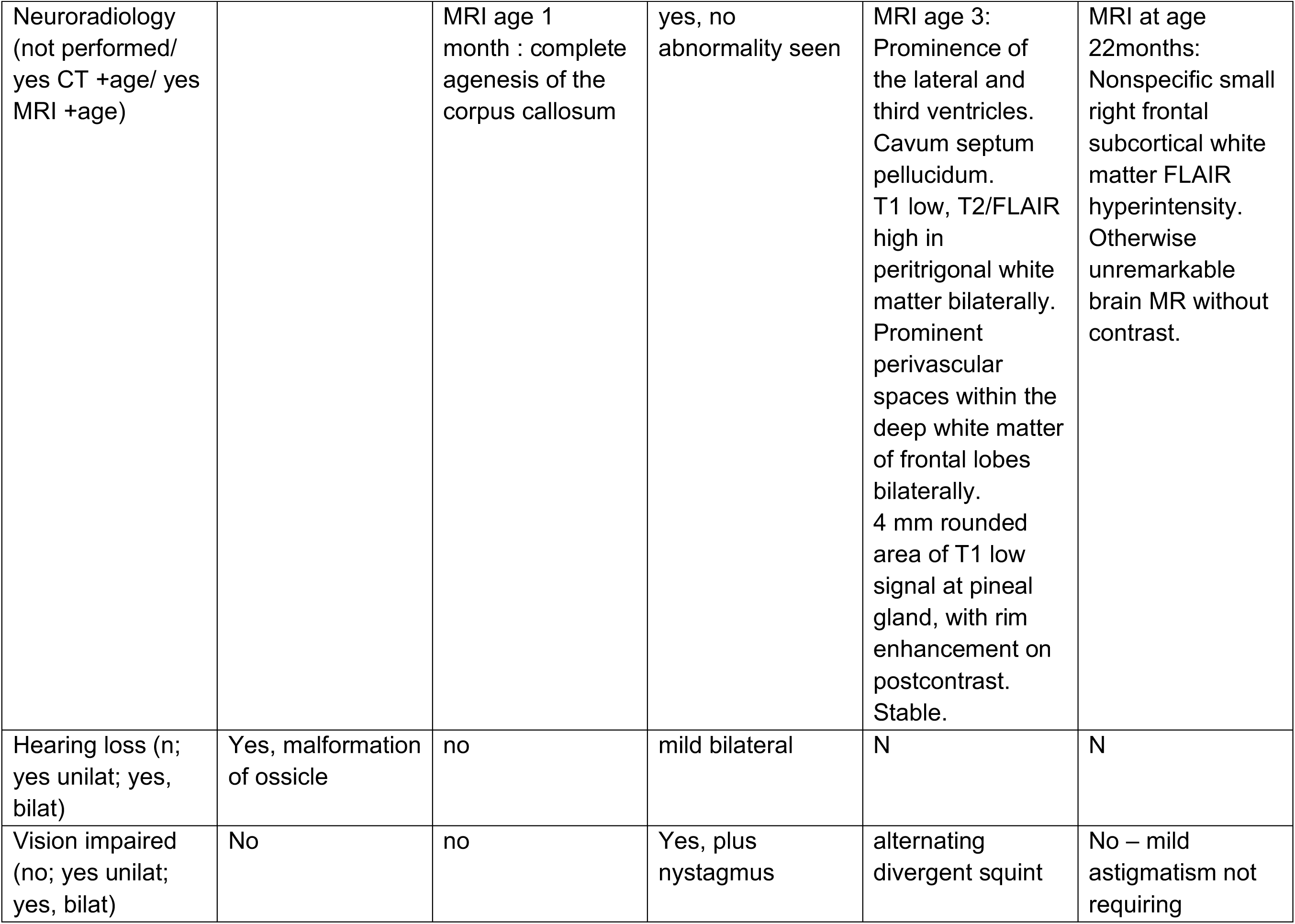
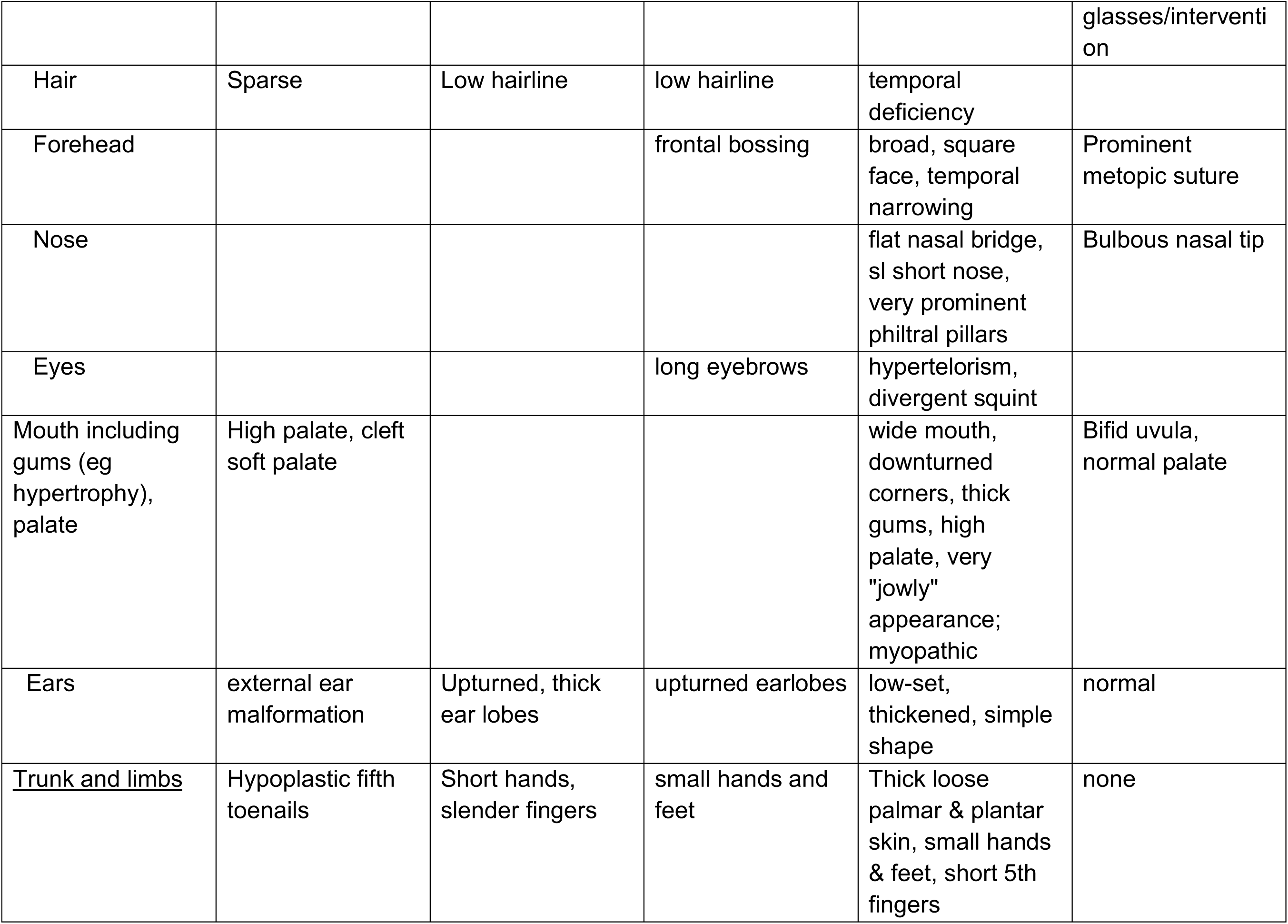
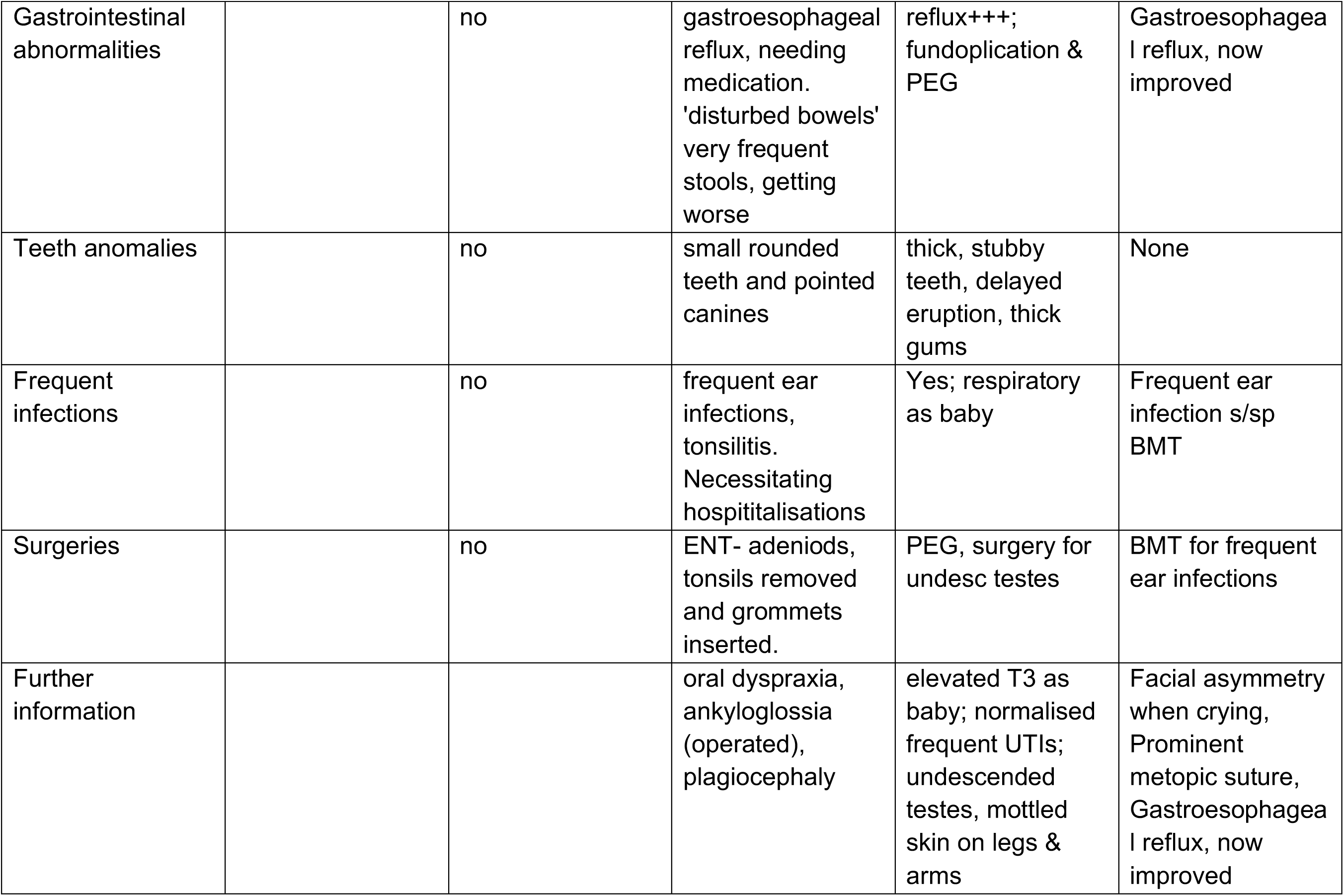
Detailed phenotypes.

**Table S4.**
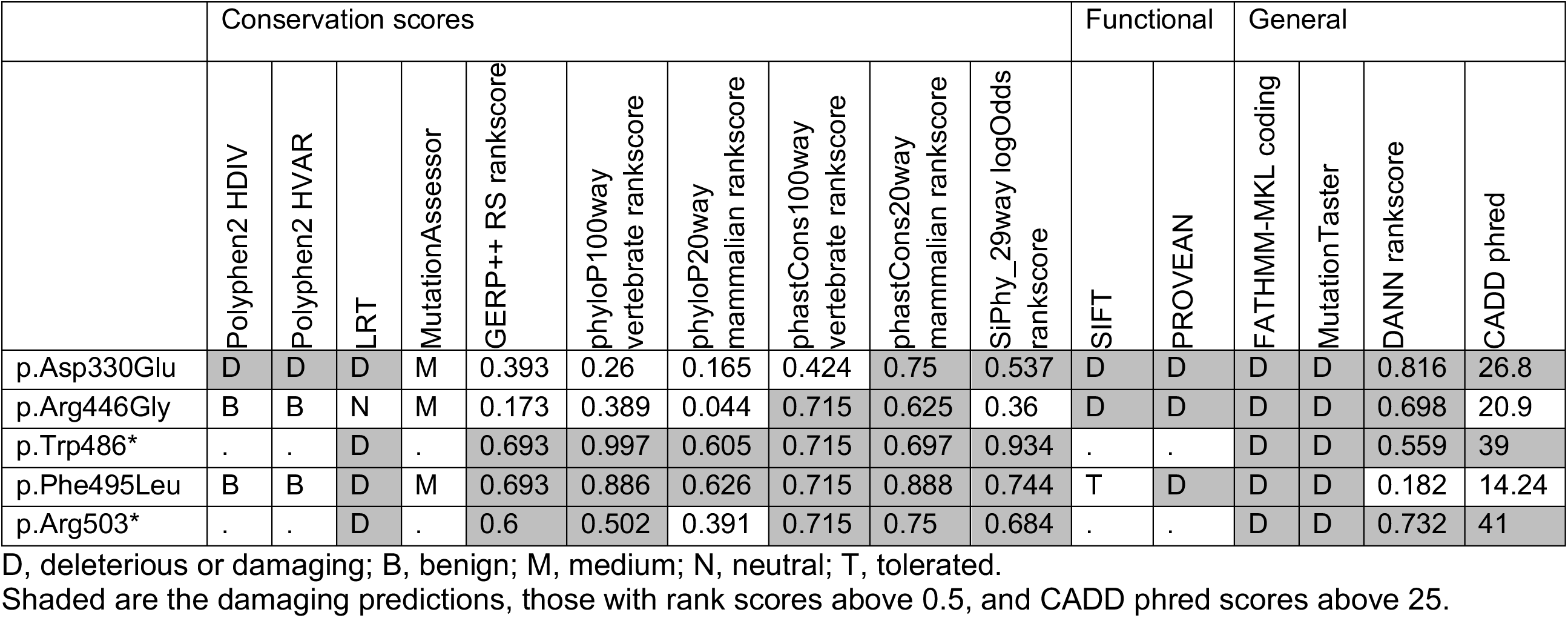
*In silico* prediction tools for the variants.

**Table S5.** Complete results comparing juvenile Bap60-KD MB with juvenile control and comparing mature Bap60-KD MB with mature control.

**Table S6.** Complete results comparing mature control with juvenile control and comparing mature Bap60-KD MB with juvenile Bap60-KD MB.

